# A genetic system for biasing the sex ratio in mice

**DOI:** 10.1101/515064

**Authors:** Ido Yosef, Liat Edry-Botzer, Rea Globus, Inbar Shlomovitz, Ariel Munitz, Motti Gerlic, Udi Qimron

**Affiliations:** Department of Clinical Microbiology and Immunology, Sackler School of Medicine, Tel Aviv University, Tel Aviv 69978, Israel

## Abstract

The ability to preselect the sex of livestock is economically beneficial and significantly increases the welfare and proper use of animals. In the poultry industry, for example, almost all males are brutally and unnecessarily killed shortly after hatching. The labor and associated costs of separation of females from males, as well as the massive killing of males, could be reduced by using a system that biases the sex of the progeny. Here, we provide a first proof of concept for such a system by crossing two genetically engineered mouse lines. The maternal line encodes a functional Cas9 protein on an autosomal chromosome, whereas the paternal line encodes guide RNAs on the Y chromosome targeting vital mouse genes. After fertilization, the presence of both the Y-encoded guide RNAs from the paternal sperm and the Cas9 protein from the maternal egg target the vital genes in males. We show that this breeding consequently self-destructs solely the males. Our results pave the way for a biased sex production of livestock, thus saving labor, costs, and eliminating substantial animal suffering.

## Main

Some aquatic organisms as well as plants that benefit from single-sex cultivation have been produced mostly by hormonal feminization of males or by masculinization of females and the subsequent production of a single-sex progeny. This was demonstrated in crustaceans^1^, fish^2-4^, and is also common in growing *Cannabis sativa*, where feminized seeds are desired (https://cannabistraininguniversity.com/feminize-marijuana-seeds). However, these practices are not feasible for terrestrial livestock.

The sex ratio in a population of mosquitoes and flies was shifted by manipulating specific genes that distort the sex ratio^5-7^. In recent breakthrough studies, researchers have even completely distorted the sex ratio, accompanied by the sterility of females, thus resulting in a collapsed population^8-12^. Such an outcome is desirable for disease-transferring insects in the wild, but not for domesticated livestock.

Sexed semen, sorted for specific gametes by flow cytometry is used for biasing the sex ratio in domesticated livestock^13^. This methodology constitutes only 5% of the artificial insemination market partially because it constitutes a substantial economic burden, shows reduced fertility rates, and its efficacy in biasing the sex ratio is variable^13^. Other approaches are therefore required.

There is currently no genetic system in mammals that biases a desired sex, while retaining a reservoir of males and females to maintain such a set-up. Manipulated animals that produce only one sex are impossible to sustain by self-crossing, because either the male or female is absent, or due to sub-fertility or infertility of the manipulated animal. Thus, despite the identification of genetic factors such as *Sry, Sox9, Foxl2*, and *Wnt4*, that determine the sex of animals, and the success of reversing animal’s sex, fertility and other genetic restrictions precluded a system that reliably produces a single sex progeny^14-18^.

We chose to provide a proof of concept in mice for an approach that produces mainly single-sex progeny while retaining a reproductive reservoir of males and females. For developing mice that produce mostly females, we used two self-sustained mouse lines, each producing males and females at an equal ratio. One line, henceforth termed the “Cas9-line”, encoded the CRISPR-Cas9 enzyme from *Streptococcus pyogenes*, expressed from a CAG promoter on an autosomal chromosome^19^. We generated the other line, henceforth termed the “Y-line”, encoding on its Y chromosome three CRISPR guide RNAs (gRNAs) targeting three autosomal genes (Fig. S1 and Appendix 1). The selected target genes, *Atp5b, Cdc20*, and *Casp8*, were all shown to be essential for mouse early development^20,21^ [*Atp5b* deficiency in mice results in embryonic lethality prior to organogenesis (https://monarchinitiative.org/gene/MGI:107801#phenotypes); *Cdc20* deficiency in mice results in metaphase arrest in two-cell stage embryos and consequently in early embryonic death^22^; *Casp8* deficiency results in necroptosis and consequently in embryonic death^23^]. We selected targeting three different genes to reduce the probability of simultaneous non-targeting of the three genes, or simultaneous in-frame corrections of these three genes, or such combinations that may result in viable males. We hypothesized that crossing these two lines would result in a progeny consisting of female-only mice, since the resulting male mice, encoding both the Cas9 and the Y chromosome gRNAs, cannot develop normally. We further hypothesized that the litter size would be half the normal size, since half of the progeny do not develop properly. To test our hypotheses, we crossed the Y-line males with the Cas9-line females, and as a control, we crossed the Y-line males with B6J females (Fig. 1a). The control cross between the Y-line males and the B6J females produced 41 pups with an average of 6.83 pups per litter (Fig. 1b). Four pups from this cross died within 3 days after birth. The cross between the Y-line males and the Cas9-line females produced 31 pups with an average of 3.88 pups per litter (Fig. 1b). Eight pups from this cross died within 3 days after birth. Physical examination of the sex of the pups at day 7 revealed a ratio of 23:14 live males to females in the cross of the Y-line males with B6J females, compared to 3:20 live males to females in the cross between the Y-line males and Cas9-line females (Fig. 1c). All mice from day 7 survived to weaning. These results demonstrate, for the first time, a genetic system for significantly biasing the sex ratio in mammals while maintaining a reservoir of fertile parents.

**Figure 1.**
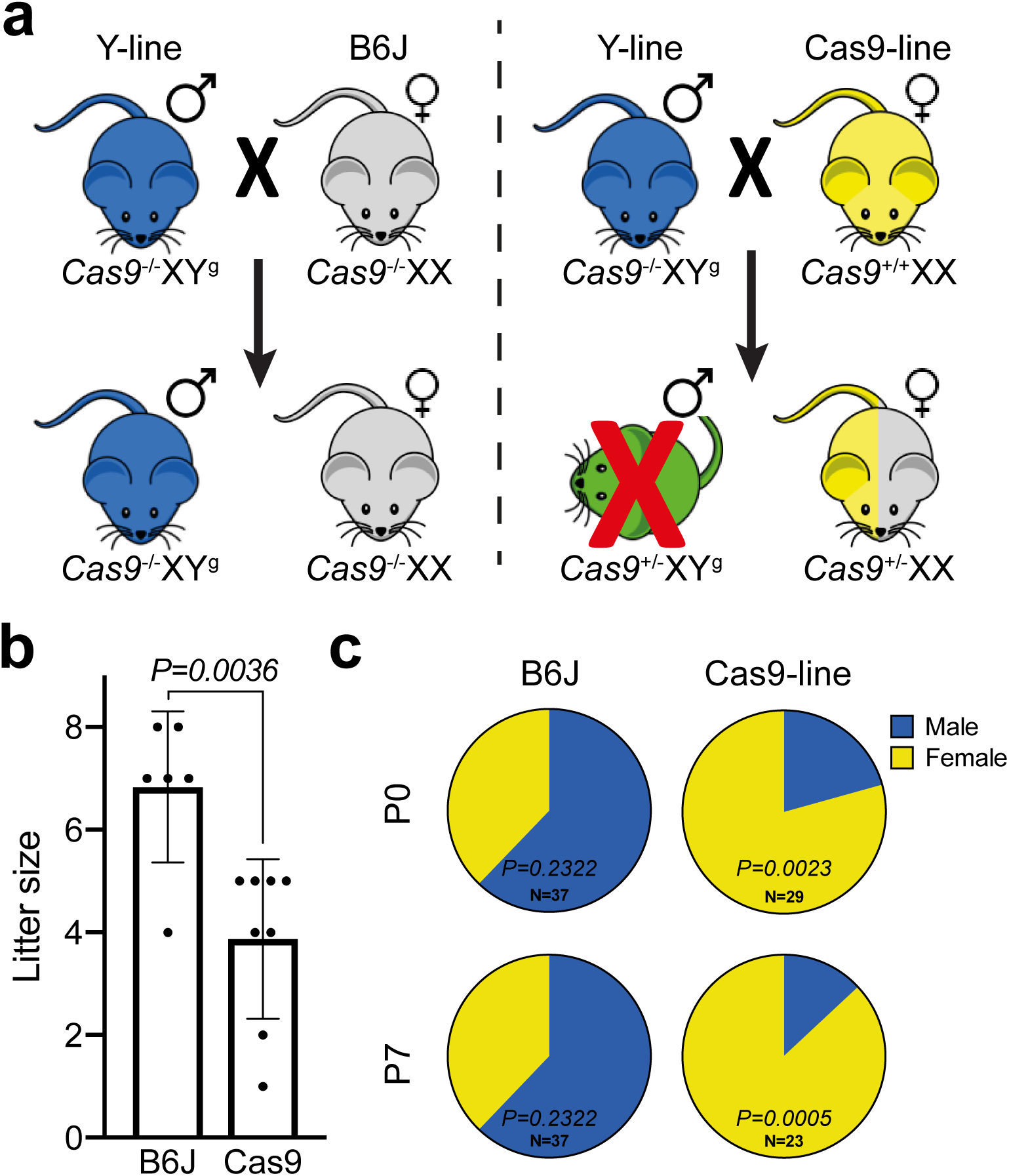
Biasing the sex ratio in mice. (a) The crosses between the Y-line males (blue) with B6J females (grey) are illustrated on the top left, and those of the Y-line males with Cas9-line females (yellow) on the top right. The presence or absence of the *Cas9* allele are annotated as *Cas9*^+^ or *Cas9*^−^, respectively, whereas the Y-linked gRNA allele is annotated as Y^g^. The X chromosome is unmodified in all instances. (b) Litter size of the crosses of the Y-line males with the indicated female line. Dots represent individual births and bars represent average ± SD. Significance was determined using a two-tailed unpaired parametric t-test. (c) Pie charts illustrating the sex distribution of the total pups from the crosses of the Y-line males with the indicated female line at day of birth (P0) and at 7 days post birth (P7). Significance was determined using a two-tailed binomial test, assuming normal 1:1 male:female ratio.

The dead mice from the cross of the Y-line with the B6J females could not be analyzed as their remains were consumed by their mother. To determine the sex of 6 pups whose remains could be distinguished out of the 8 dead pups from the crosses between the Y-line males with the Cas9-line females, we carried out PCR amplifying the Y chromosome. These analyses revealed 3 males and 3 females among these 6 pups. One of these males was deformed, lacking developed limbs, and most likely died upon birth (Fig. S2a) suggesting that the genetic system eliminated this male. The other two males appeared smaller compared to their siblings (Fig. S2b,c) suggesting that their premature death is also due to the genetic system. Indeed, DNA sequencing of the three target genes demonstrated that two or three genes were disrupted in two out of two analyzed males (Fig. S3). We further analyzed DNA from one of the surviving male from the cross between the Y-Line males and Cas9-line females. In this case, two genes were disrupted (Fig. S3). These results indicate that all tested males were targeted, but also that lethality could be delayed or abolished probably due to differences in the type and extent of disruption of the target genes. These sporadic occurrences of male late lethality or even viability could probably be eliminated by simultaneous targeting of more genes using gRNAs that target multiple chromosomal regions^24^ or by addition of gRNAs targeting more genes/regions than the current system.

Based on similar principles, one can also establish lines producing only male progeny. For such an outcome, the paternal line should be engineered to encode the gRNAs on its X chromosome, and should be crossed with the maternal Cas9-line (Fig. S4a). Applying this system in animals in which the female is the heterogametic organism, such as chicken, where the female carries the Z and W sex chromosome, and the male carries two copies of the Z chromosome^25^ can also be manipulated similarly. In these cases, for female-only progeny, the Z chromosome of the maternal line should encode the gRNAs, and the paternal line should encode Cas9 (Fig. S4b). For male-only progeny, the maternal W chromosome should encode the gRNAs and paternal line should encode Cas9 (Fig. S4c). Thus, this system can easily be manipulated to accommodate changing requirements in different organisms and for both genders.

The females obtained using this approach are genetically modified organisms (GMO) because they retain the Cas9 enzyme in their genome. In mice, these are normal-looking, fertile, and viable animals^19^. Nevertheless, from a regulatory point of view, a non-enzymatic transgene, such as short gRNAs, may be considered more tolerable than the Cas9 nuclease is. Thus, the system can be modified to obtain females that instead encode the gRNAs in their genome, if in that case Cas9 is encoded on the paternal Y chromosome. In this respect, it is noteworthy that a GMO salmon, which grows faster than its parental non-GMO strain, owing to a transgene regulating a growth hormone, has been approved by the US Food and Drug Administration for the food industry^26^. Thus, GMOs may, in principle, be approved for food production.

Using the system shown here in the poultry industry has the potential for an annual saving of approximately seven billion newborn male chicks from cruel death by either suffocation or live-grinding^27^. This use will require minimal manual sexing, as male eggs will be significantly reduced or even eliminated with an optimized system, thus saving also costs and labor. Additionally, it may be used for producing a single-sex population of cattle, swine, desired fish, crustaceans, as well as feminized seeds of certain crops such as *Cannabis sativa* and *Humulus lupulus*. Thus, apart from animal welfare and ethical considerations, the use of the system will unleash huge economic benefits and savings.

## Methods

### Mice

All mice were bred under specific pathogen-free conditions in the animal facility at Tel Aviv University. Experiments were performed according to the guidelines of the Institute’s Animal Ethics Committee.

Mice of the Cas9-line were purchased from Jackson laboratories (Stock No: 026179; Rosa26-Cas9 knockin on B6J)^19^. These mice encode a cassette in the Rosa26 locus on chromosome 6 constitutively expressing the *Sp*Cas9 endonuclease from a CAG promoter.

Mice of the Y-line were constructed by Cyagen Biosciences (California, USA). These C57BL/6N mice encode the following guide RNAs on their Y chromosome: 5’-CACTGCCACCGGGCGAATCG-3’; 5’-CAGACCTGAATCTTGTAGAT-3’; 5’-TGCAGAGATGAGCCTCAAAA-3’ targeting the genes *Atp5b, Cdc20*, and *Casp8*, respectively. These guides were cloned into a vector targeting the reverse orientation of the 2^nd^ exon of the Y chromosome *Uty* gene, which is not part of the pseudoautosomal Y region. Figure S1 provides a schematic summary and Appendix 1 provides detailed description of the Y-line construction.

### Crosses and sex determination

Males from the Y-line were crossed with females from the Cas9-line and with B6J females. Sex was determined by observing the genitals at day 7 or by PCR of the Y chromosome on DNA extracted from the animal’s tissue. Sanger sequencing of the target regions was carried out following PCR amplification of these regions (see Tables S1+S2 for oligonucleotides and PCR set-ups).

### Statistics

Data are presented as the mean ± SD. Comparisons were performed using two-tailed unpaired parametric t-test or two-tailed binomial test, assuming normal distribution or two-tailed t-test of the variance–covariance matrix of the standard errors.

## Data Availability Statement

The data that support the findings of this study are available from the corresponding authors upon request.

## Acknowledgements

The research was funded by the European Research Council StG program (grant 336079), the European Research Council PoC program (grant 811322), the Israel Science Foundation (grants 268/14, 1416/15), the Israeli Ministry of Science (grant 3-14351), and the Varda and Boaz Dotan Research Center.

## Author Contributions

Conceptualization, I.Y., U.Q.; Methodology, I.Y., L.E.B., R.G., I.S., A.M., M.G., and U.Q.; Investigation, I.Y., I.S., R.G., L.E.B., M.G., and U.Q.; Writing – Original Draft, U.Q.; Writing – Review & Editing, I.Y., M.G., A.M., and U.Q.; Funding Acquisition, M.G. and U.Q.; Supervision, M.G. and U.Q.

## Supplementary Material

**Figure S1.**
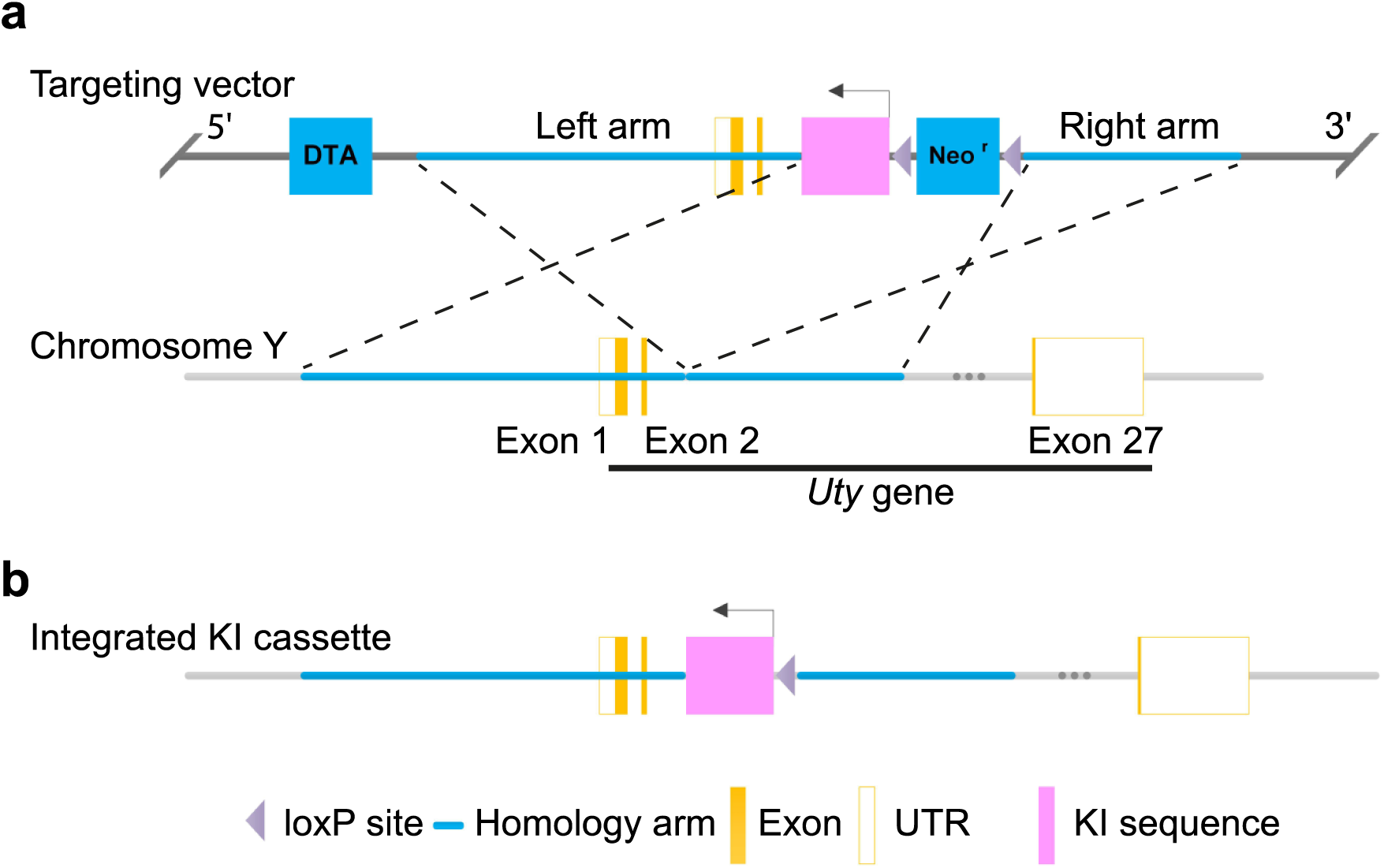
Schematic summary of Y-line generation. (a) Mouse genomic fragments containing homology arms to the *Uty* gene located on mouse chromosome Y were amplified from a BAC clone, and sequentially assembled into a targeting vector along with negative and positive selection markers (DTA and Neo, respectively). The KI cassette, encoding the gRNAs targeting genes *Atp5b, Cdc20*, and *Casp8* was inserted into the targeting vector, which was designed to integrate in reverse orientation of the 2^nd^ exon of the *Uty* gene. (b) The KI cassette integrated into the *Uty* gene and the Neo cassette was self-deleted due to the loxP sites. Details on the linearization of the targeting vector, its transfection into C57BL/6N ES cells, as well as PCR and Southern blot analyses and ES implantation procedures are provided in Appendix 1.

**Figure S2.**
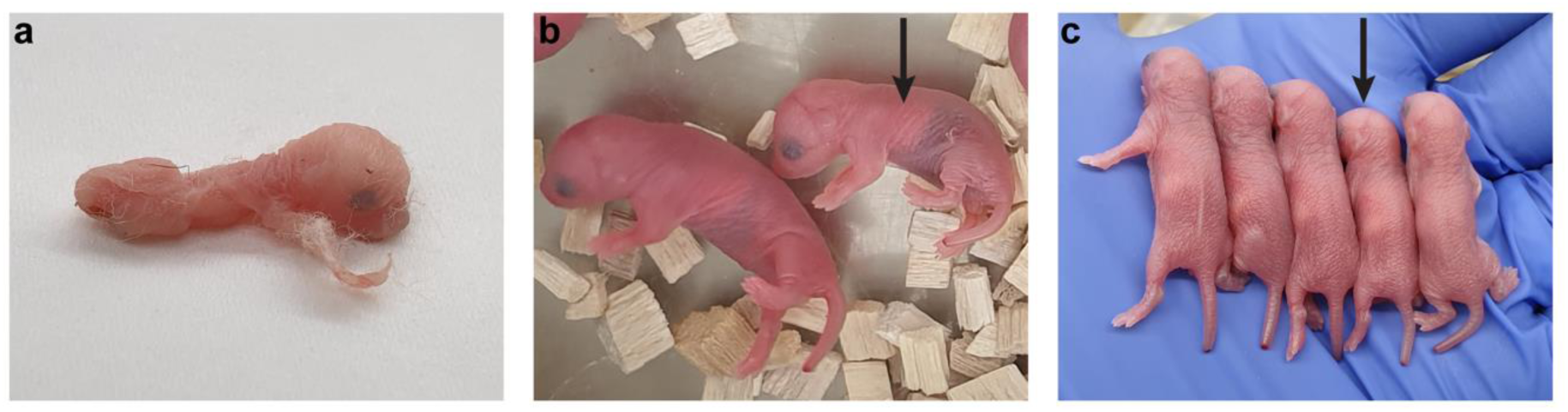
Pictures of the (a) deformed and of (b,c) smaller (arrowed) males born from a cross between the Y-line males and the Cas9-line females.

**Figure S3.**
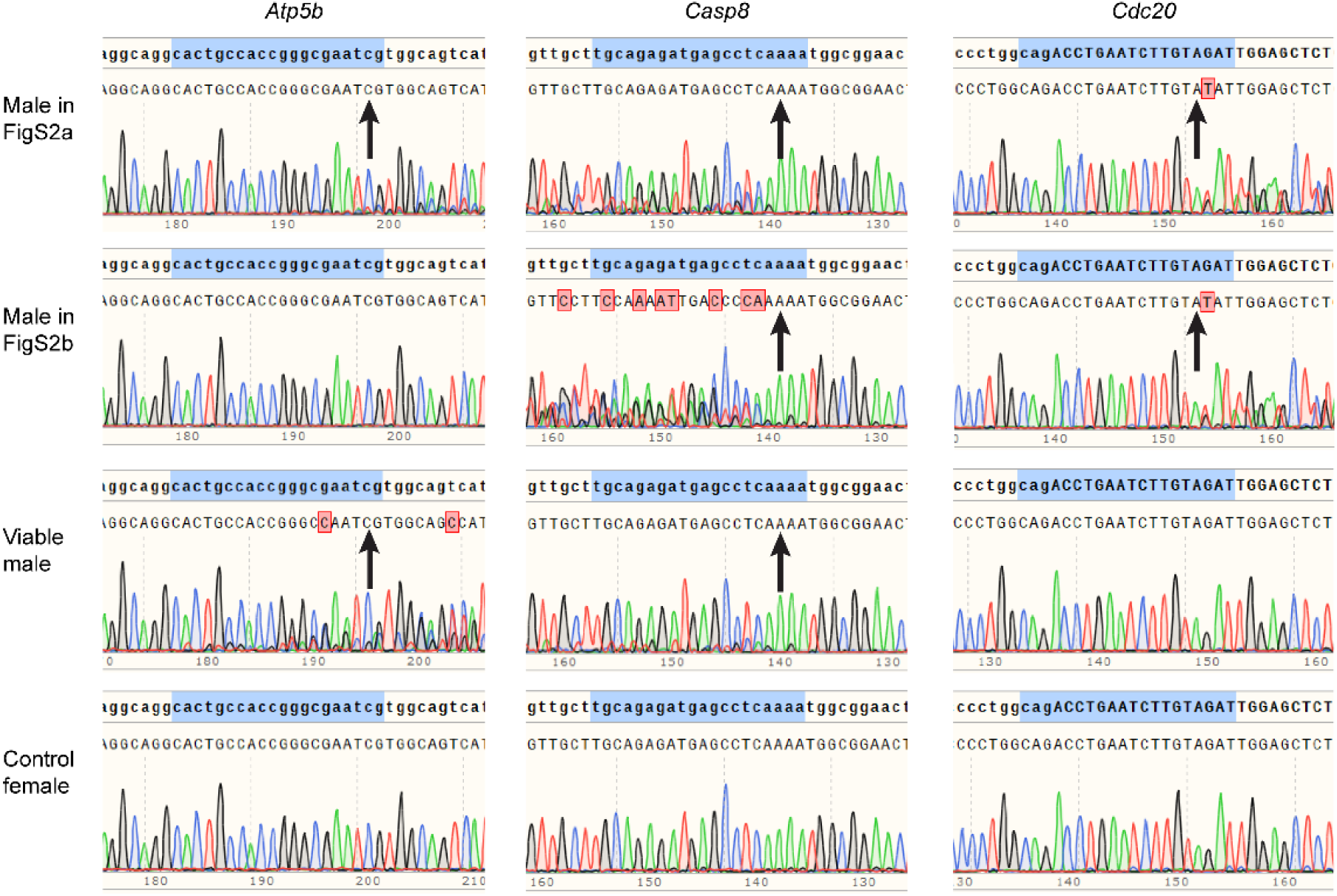
Chromatograms of Sanger DNA sequencing of the PCR amplified genes *Atp5b, Casp8*, and *Cdc20*, from samples of the indicated dead males, a viable male, and from a control normal dead female. Targeted regions homologous to the gRNAs are highlighted in blue. All samples were taken at P0 from pups of the Y-line males and the Cas9-line females cross. Arrows point to indel locations (*P<0.001*, two-tailed t-test of the variance–covariance matrix of the standard errors), as determined by Tracking of Indels by Decomposition^28^.

**Figure S4.**
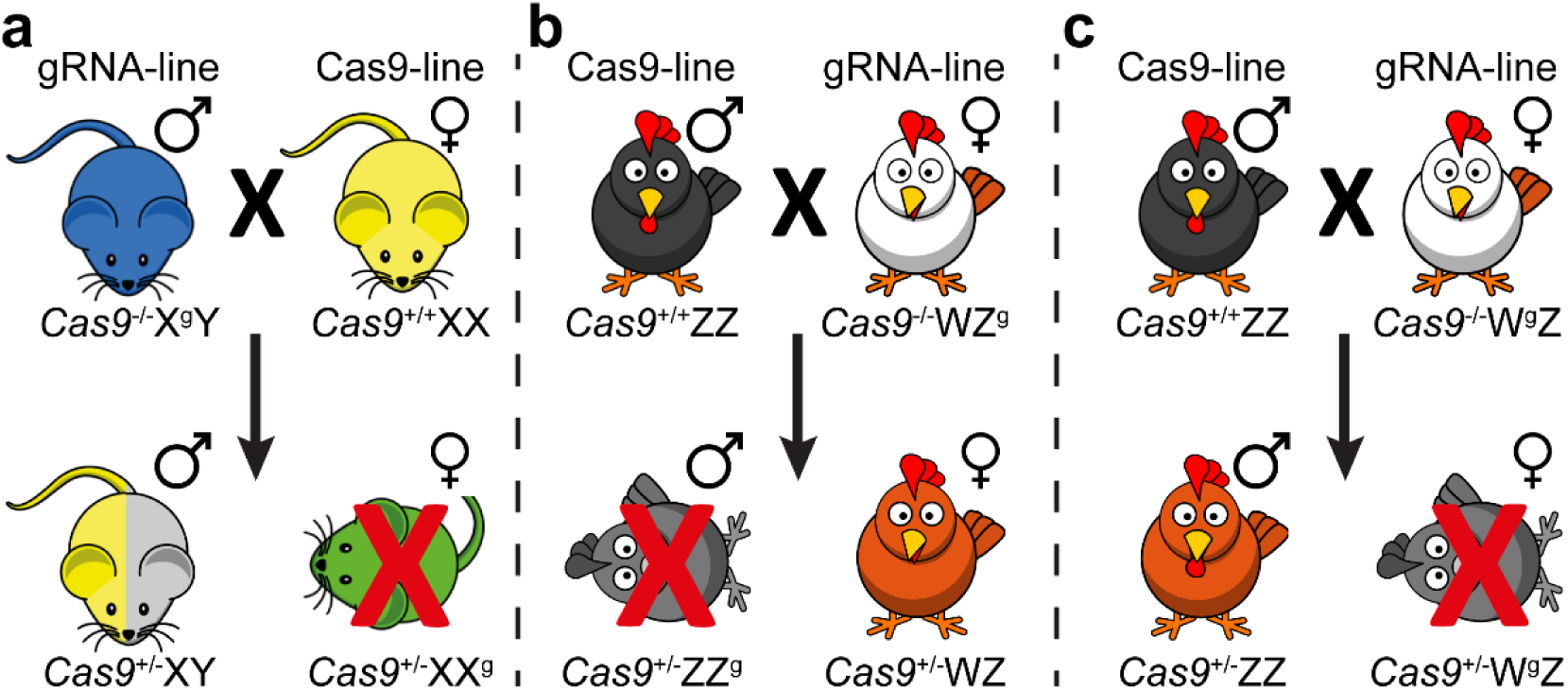
Schematic illustrations of hypothetical crosses between (a) mice producing only males (b) birds producing only females (c) birds producing only males.

**Table S1.**
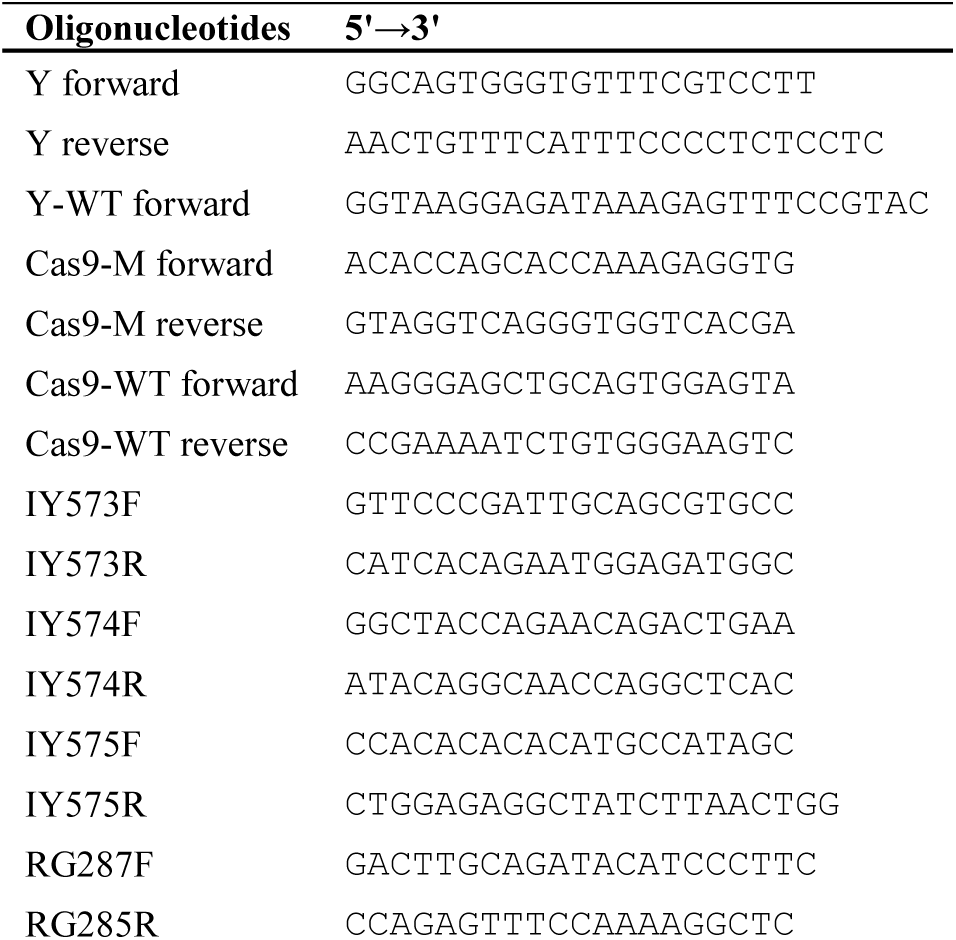
Oligonucleotides used in this study

**Table S2.**
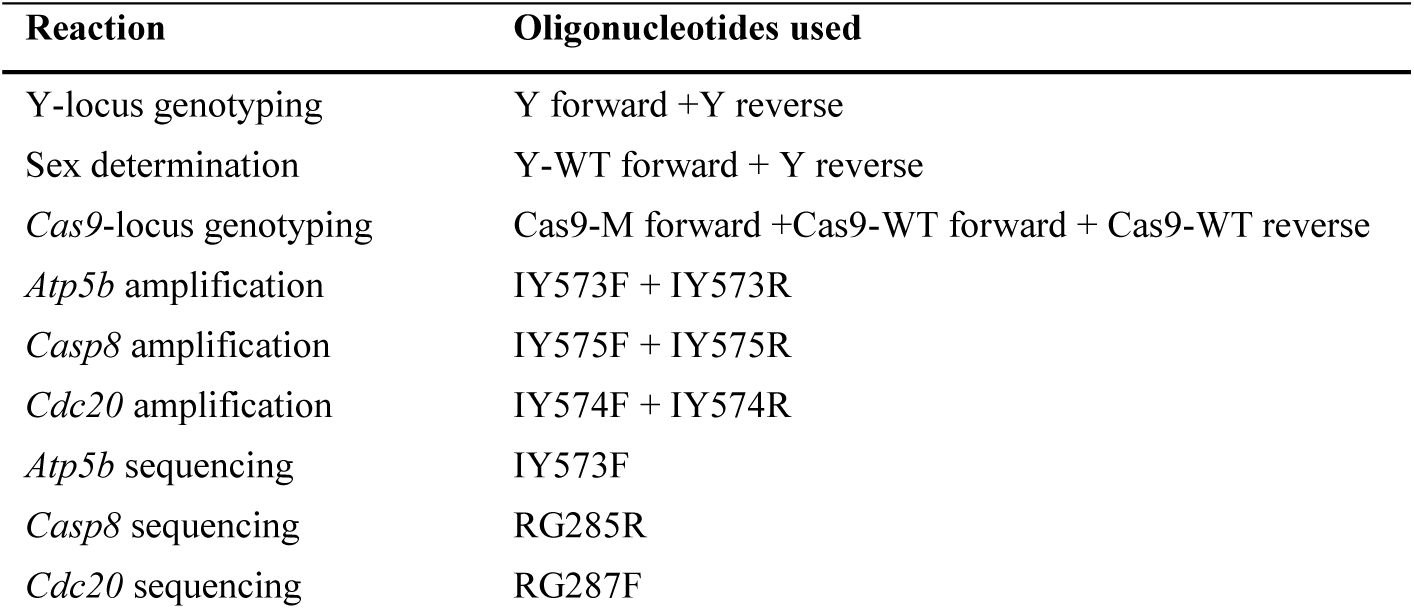
PCR set-ups

